# Ultrasound inhibits SARS-CoV-2 infectivity in vitro

**DOI:** 10.1101/2022.11.21.517338

**Authors:** Flávio P. Veras, Gilberto Nakamura, Marcelo A. Pereira-da-Silva, Ronaldo Martins, Eurico Arruda, Fernando Q. Cunha, Odemir M. Bruno

**Affiliations:** Institute of Biomedical Sciences, Federal University of Alfenas, Minas Gerais 37130-001, Brazil; São Carlos Institute of Physics, University of São Paulo, São Carlos 13566-590, Brazil; Department of Clinical Analyses, Toxicology and Food Science (DACTB) School of Pharmaceutical Sciences of Ribeirão Preto, University of São Paulo, Ribeirão Preto 14049-900, Brazil; Center of Research in Inflammatory Diseases, Ribeirão Preto Medical School, University of São Paulo, Ribeirão Preto 14049-900, Brazil

## Abstract

The global spreading of SARS-CoV-2 and the emergence of new variants underscore the ongoing need to develop new vaccines and antiviral drugs. While electromagnetic and acoustic waves have well-known virucidal properties, their application in therapeutic settings has been limited due to harming effects in biological matter. Here, we investigates the potential of ultrasound to interfere with SARS-CoV-2. Specifically, we study the effects of acoustic waves in the 1–20 MHz frequency range to determine their impact on the viral envelope of SARS-CoV-2. Our in vitro experiments demonstrate ultrasound exhibits a virucidal effect on SARS-CoV-2 without production of heat or cavitation. This study offers a promising physics-based approach to combat SARS-CoV-2 and potentially other spherical viruses, broadening the scope of antiviral treatments.

## I. INTRODUCTION

Coronaviruses are positive single-strand RNA viruses with a characteristic crown-like shape responsible for acute respiratory and enteric diseases afflicting mammals, birds, and humans^1^. SARS-CoV-2 caused the 2020-2021 pandemic, which reduced global life expectancy by 1.8 years with an estimated burden of over 12 millions estimated deaths^2^. Although vaccination remains the primary disease prevention and control method, the continued global spread of SARS-CoV-2 has promoted the emergence of new variants with significant mutations in the spike protein. These mutations often change the antigenic identity of the virus, and potentially reduce the immuno-response against strains not covered by vaccines. Therefore, there is an urgent necessity to elaborate new methods to complement the development of vaccines and antiviral drugs.

While biological and chemical approaches have been the focus of most research and disease control strategies, there is a significant gap for physics-based options. Ionizing radiation – short electromagnetic waves in the ultraviolet frequency range and above – interact strongly with electron-rich regions, including nucleic acids, resulting in a fast and cheap virus inactivation method. However, they can also damage cells and their components, including DNA/RNA, and thus find little *therapeutic potential for ongoing viral infections*. Non-ionizing radiation can also damage biological tissues. For instance, ultrasonic acoustic waves can resonate with bubbles in solution (cavitation) in certain circumstances, leading to mechanical damage after bubble collapse, or chemical damage during expansion-contraction cycles due to the production of free radicals, including hydroxyl. Despite the grim scenario, the likelihood of cavitation events decrease with higher frequencies thus opening a venue for therapeutic applications.

A promising strategy employs acoustic waves to resonate with the viral structural proteins, including the spike protein. The numerical study by Wierzbicki suggested that acoustic waves with frequencies between 100-500 MHz could resonate with the SARS-CoV-2 envelope and associated proteins, thereby neutralizing it^3^. In a revised study, the authors show ultrasound between 1 and 20 MHz could damage specific parts of spike proteins^4^.

In this work, we carry out in vitro experiments to verify whether SARS-CoV-2 can be inactivated by acoustic waves, and the corresponding effects over the viral envelop, if any. The frequency range 1–20 MHz falls in the same range of ultrasound devices for medical imaging, which are, in practical applications, safe for therapeutic uses. Although the studies^3,4^ alludes to ultrasound-produced resonances, this effect has not yet been experimentally proven. Our findings indicate that acoustic waves can produce a virucidal effect, without any noticeable change in temperature or pH. Our working hypothesis is that the mechanical properties and the spherical geometry of SARS-CoV-2 play a crucial role in the energy transfer from the ultrasound to the virus.

This approach offers a unique method to combat SARS-CoV-2, and can be explorer for other spherical viruses, expanding our arsenal of antiviral treatments. The paper goes as follow. Section II presents our main results, detailing the observed effects of ultrasound on the virus. In Section III, we discuss the implications of our findings, with concluding remarks in Section IV. Section V details our experimental setup, virus strains, and methods.

## II. RESULTS

We evaluated the potential virucidal effects of ultrasound on stocks of SARS-CoV-2 using ultrasound imaging devices for medical diagnostics. Our choice to use imaging devices instead of piezoelectric transducers was twofold. First, the frequencies for imaging devices lies in the 1–20 MHz range. Second, in addition to their safety protocols in place, their power output is generally low to ensure patient safety. Modern devices also indicate the mechanical index, MI ∝ *p/f* ^1*/*2^, an index that indicates the likelihood of cavitation events for pressure *p* and frequency *f*. As a general guideline, MI values below 0.3 are cavitation free (low signal and high frequency), between 0.7–1 can support non-inertial cavitation (bubble oscillations) with moderate probability, and MI *>* 1 support both inertial and non-inertial modes with high probability. In our experiments, MI was kept between 0.3 and 0.5, below the suggested 0.7 threshold. Taken together, the results indicate that imaging devices are suitable candidates sources of ultrasound for therapeutic applications.

We used virus stocks for Wuhan, Delta, and Gamma variants of SARS-CoV-2 in order to investigate whether the virus-ultrasound interaction would affect conserved regions of the viral structure or would be variant-dependent. The solutions containing SARS-CoV-2 were exposed to three ultrasound transducers for 30 minutes (Fig. 1A). Since each transducer can produce various frequency profiles, we aggregated the results over several modes as the representative data for the transducer (see below for specific frequency dependence). For the sake of simplicity, we label the equipment as Equip. A (Philips; Envisor HD; linear transducer 3–12 MHz), Equip. B (Esaote; MyLab 60; linear transducer 5–10 MHz), and Equip. C (Esaote; MyLab 60; linear transducer 6–18 MHz). We included two control groups for each strain: a primary control consisting of the untreated viral solution, and a secondary control sample containing no virions (mock). We confirmed the effectiveness of infections by the primary control group by comparing the incubation of both control samples in Vero-E6 cells (Fig. 1B top). Infection of Vero-E6 cells with US-treated samples were analyzed 24 hours post-infection (hpi) via immunofluorescence and confocal microscopy to determine the extent of the virus infection (Fig. 1B). This was done by staining the SARS-CoV-2 spike protein and double-stranded RNA (dsRNA) to gather evidence on the viral replication process. Virucidal effects were optimal across all strains for the transducer with bandwidth 5–10, centered at 7.5 MHz. The virucidal effect was reduced for the remaining devices for all but the Wuhan strain.

**FIG. 1.**
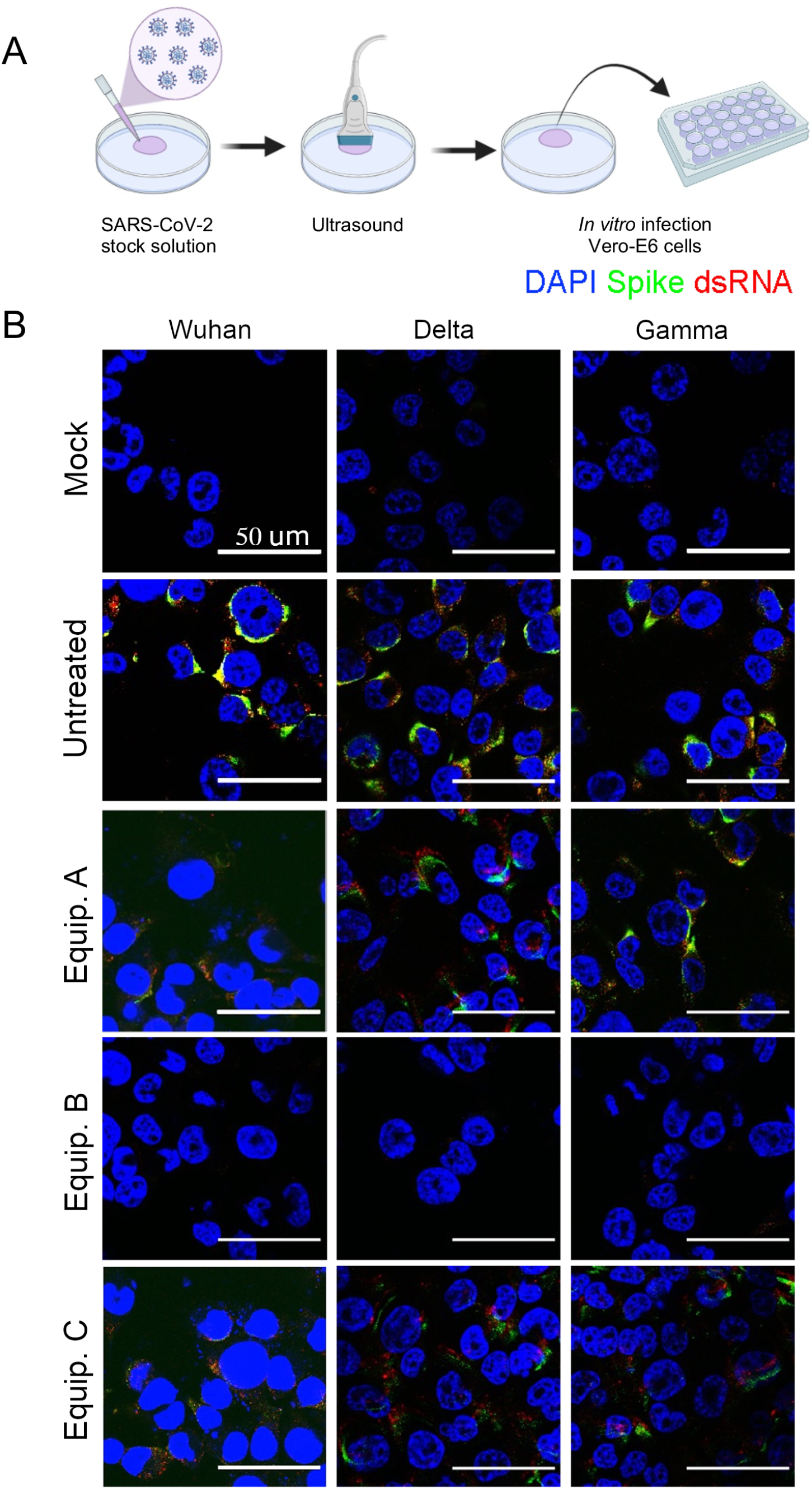
US treatment inhibits SARS-CoV-2 infection and replication. (A) Representative model of US exposure. (B) Immunofluorescence analysis of Spike (green) and dsRNA (red) expression of SARS-CoV-2-infected Vero-E6 cells after ultrasound exposure. DAPI (blue) was used for nuclei staining. Scale bar indicates 50 *μ*m. Data are representative of at least two independent experiments.

To complement and quantify our initial findings, we further explored how the ultrasound frequency impacts virus infectivity (Wuhan) after the 24 hpi mark by conducting TCID50 titer assay at various frequency profiles. For this experiment, we used the linear transducer L12-3 (Philips, 3–12 MHz with 38 mm frontal aperture). The Wuhan strain was exposed to the transducer at various frequency profiles, labeled by the center of the mode, as shown in Fig. 2, during 1, 5, and 10 minutes. Consistent with our previous findings, modes centered around 7.5 MHz are far more effective imparting permanent transformations in viral proteins associated with host infection, regardless of exposure time. Fig. 2A reveals lower frequencies are less effective. This result suggests that cavitation cannot be the main driver of the effect because it would be in contradiction with the increased chance of cavitation events, which scale as 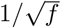. At the same time, increased exposure decreases TCID50 across all modes. Our working hypothesis is that the other frequency profiles contains the resonant frequency around 7.5 MHz albeit at lower percentage, resulting in slower energy deposition rates.

**FIG. 2.**
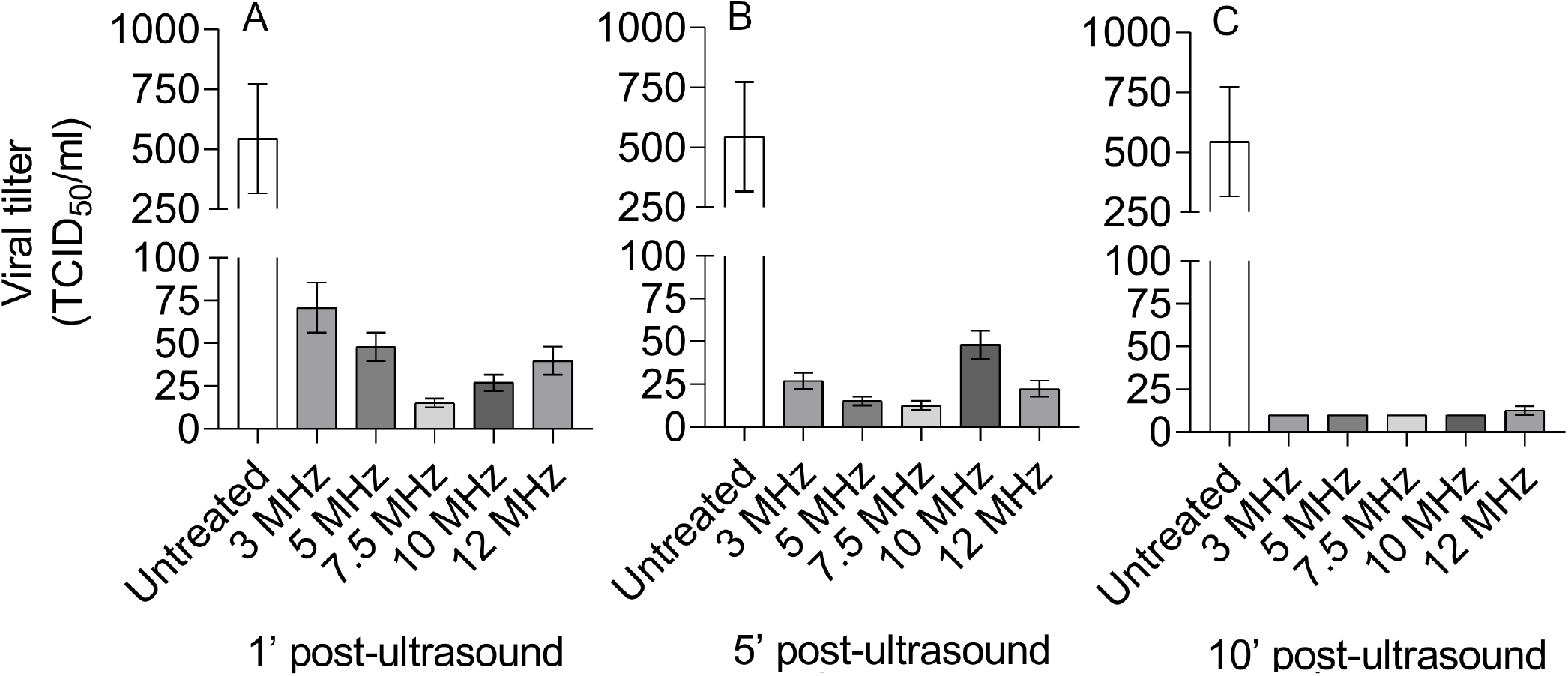
Efficiency of ultrasound treatment depends on the frequency. TCID50 essay for the SARS-CoV-2 Wuhan strain after ultrasound exposure during 1, 5, and 10 minutes at various frequency profiles in Equipment A. Each profile is labeled by the dominant frequency.

Another significant factor that can interfere with viral infectivity is denaturation of the viral capsid or spike protein induced by high temperatures^5^. Acoustic radiation can deposit energy in biological material and thus increase its temperature due to the increased displacement of molecules during compression-rarefaction cycles. We monitored the temperature of culture medium during the ultrasound treatment. No appreciable temperature increase was observed that could denaturation viral proteins (Fig. 3).

**FIG. 3.**
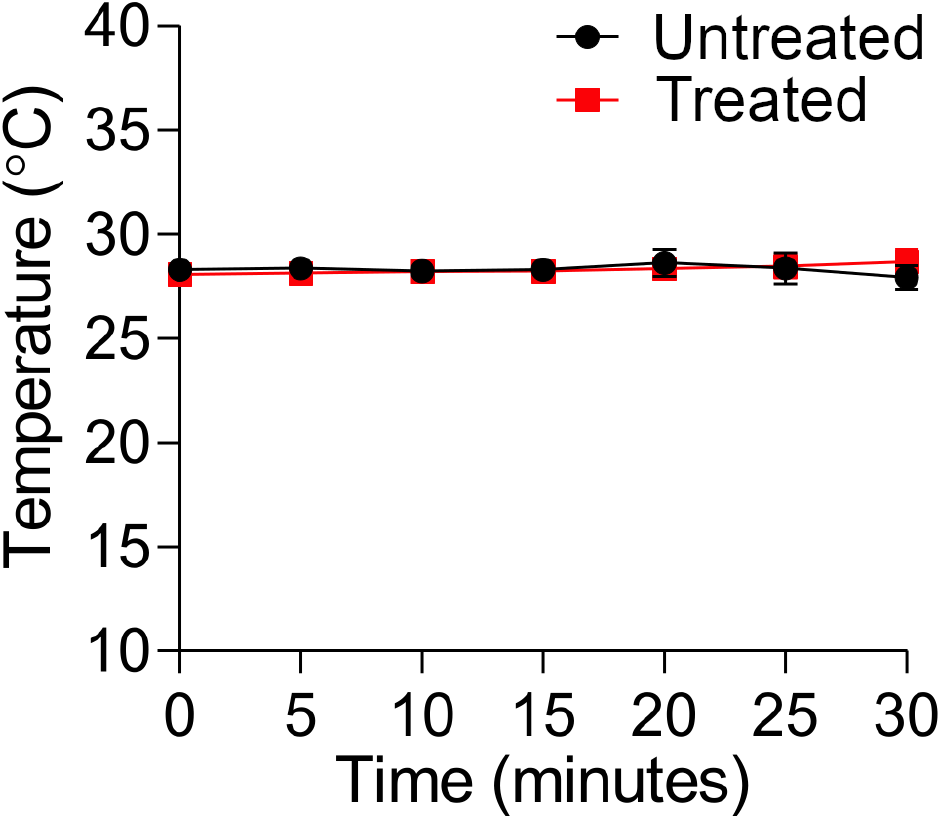
Ultrasound exposure did not alter medium culture temperature. Quantification of DMEM medium culture temperature by a thermal camera for 30 min after exposure by Equipment A with linear transducer 3–12 MHz.

We used scanning electron (SEM) and atomic force microscopy (AFM) to assess whether ultrasonic waves alter the morphology of SARS-CoV-2. The SEM images (Sigma Zeiss FE-SEM, Carl Zeiss) for control and treated samples (Wuhan) were taken at 2 kV. The control group exhibit spherical objects between 80–100 nm, either isolated (Fig. 4A) or in aggregates. The microscope ‘s resolution was insufficient to capture finer details such as spike proteins due to operational limitations. The morphology of the virus in the treated group, however, had defective surfaces upon closer inspection and smaller diameters when compared to the control group (Fig. 4B inset). The line profile from the treated particles indicates the original spherical surface transformed into a significantly rougher texture, characterized by prominent depressions and valleys.

**FIG. 4.**
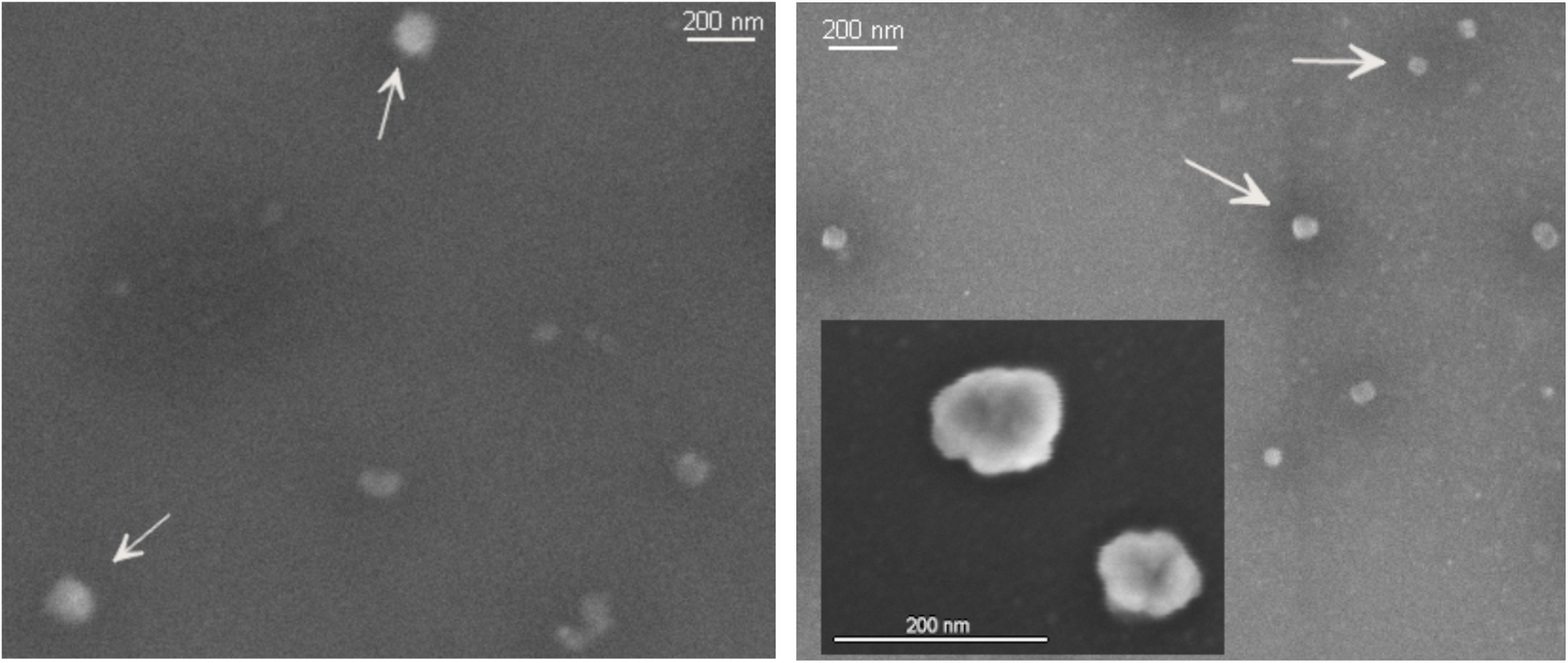
Effects of US treatment on SARS-CoV-2/Wuhan. The morphology of the control sample (left) shows notable differences from UV-inactivated viral particles after US treatment (right). SEM images produced at 2 kV.

A direct assessment of the envelope was done via AFM in tapping mode (see Fig. 5). Besides the topology, AFM can also map the spatial distribution of defects and mechanical properties of the target surface via the phase contrast, with little to no damage to the biological sample. The images reveal that treated particles breaks apart, reducing in size when compared to the control group. More importantly, the nearly spherical surface dotted with slight elastic elevations (spikes proteins, creating a thin noisy layer over the spherical centers), becomes rigid and fragmented, suggesting absence of spikes and ruptured envelope (see Fig. 5 top). To shine some light on the issue, a color correction was employed to highlight only the upper mid-height of the structures (Fig. 5 bottom). The procedure demonstrates that the envelope breaks apart, resembling a popcorn. Unfortunately, it remains unclear how the envelope breaks and how to reconstruct them from the AFM images alone.

**FIG. 5.**
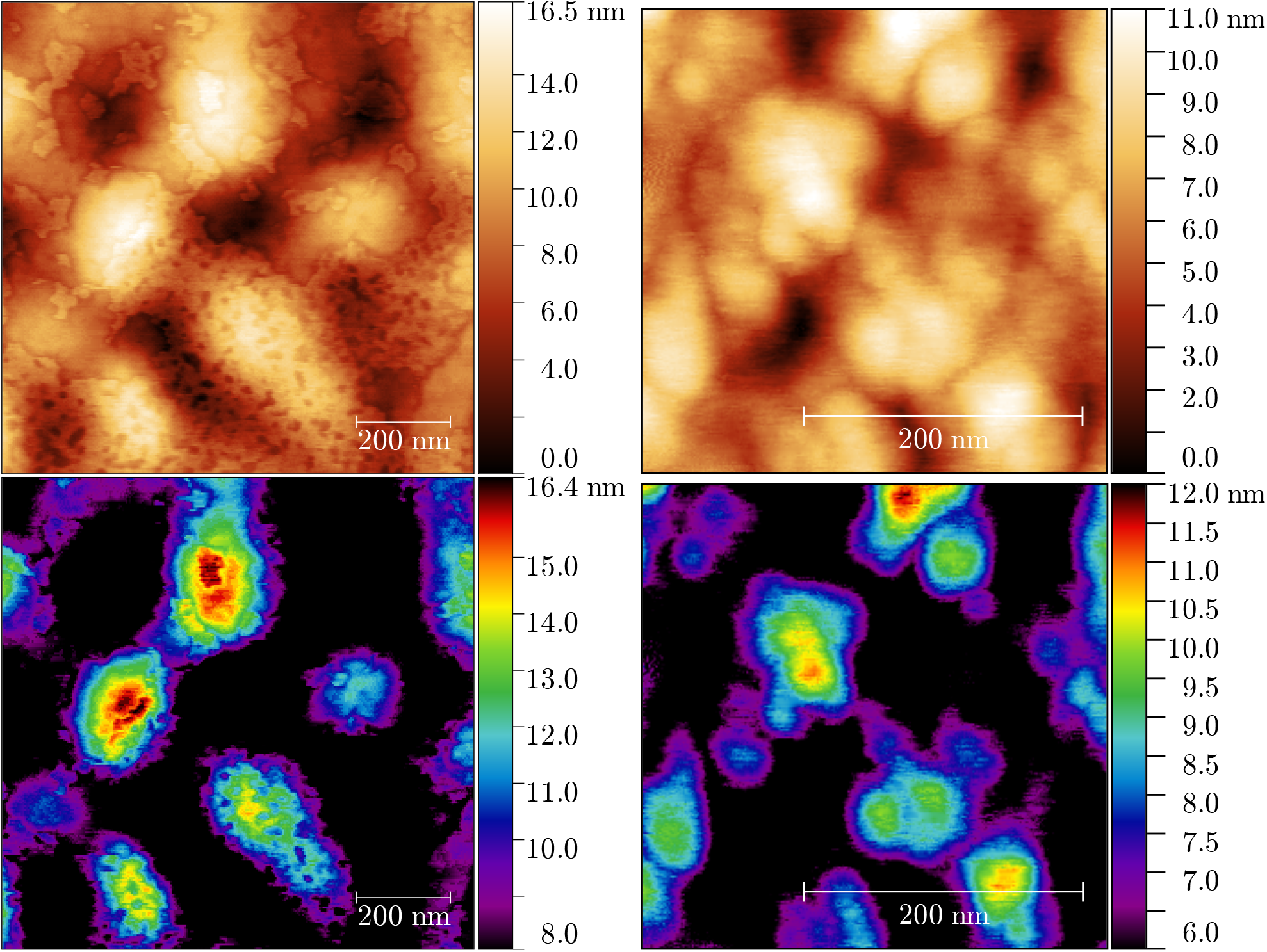
Viral morphology after exposure to US waves, SARS-CoV-2/Wuhan. (left) Height distribution for control group at (top) full and (bottom) half-height. (right) US acoustic waves disrupt the integrity of the viral shell. AFM images produced in tapping mode.

## III. DISCUSSION

### Advances on antiviral therapies

Over the past 30 years, over a hundred new antivirals have been approved by the Food and Drug Administration, about half of which are HIV-related treatments^6^. Formally, antivirals therapies inhibit the replication cycle by blocking or disrupting key steps in the virus cycle, including membrane adsorption or genome replication. Proposals of new candidate molecules require a precise understanding of the mechanisms describing the viral activity in the host cell. Even more, because virus appear in a variety of forms – with a plethora of binding proteins and specialized enzymes – and they hijack the host machinery for their own metabolism, antivirals must be highly specific to prevent further harm done to the host environment. For example, the HIV antiviral enfuvirtide binds to the glycoprotein gp-41, interfering with the virus adsorption and penetration into the membrane without disturbing the expression or binding to unintended host proteins.

Although high specificity is often desirable, it also may create a bottleneck when dealing with different virus strains. New candidate molecules target structures shared by a broader spectrum of virus, such as Remdesivir, an antiviral that acts on viral RNA polymerases of MERS and SARS coronaviruses. A similar effect can also be achieved by modulating the immune response of the host, via interferons, to disrupt the viral replication inside the cell. Interferons are innate components of the immune system that trigger the expression of antiviral proteins such as interferes with the virus, either on infected or healthy cells, such as ribonucleases – enzymes that degrade small RNA, including vRNA.

Alternative antiviral technologies that rely on physical rather than biomolecular signatures have been of particular interest to complement current therapies. In particular, nanoparticles^7^. These structure often have metallic or polymeric cores and are coated with membrane-like components to increase bio-compatibility with the target, resembling virus-like particles. They have been employed in a multitude of medical applications, including disease diagnosis, cancer treatments, and as drug delivery vectors, ie, they protect the drug during the transport and release it at the target, activated by electromagnetic or mechanical fields. Nanoparticles can also be coated with antibodies to degrade viral capsids directly in localized regions of the body.

Despite the variety of antiviral therapies available, a broad spectrum solution remains elusive, with specificity and delivery mechanism being the main culprits. In addition, certain tissue can pose additional challenges due to selective permeability. This tends to occur more often near interfaces between tissues, such as the blood-brain barrier for example, and greatly hinders the effectiveness of antivirals. The issues could be addressed by using radiating electromagnetic or acoustic fields. Radiation fields find many applications in medical applications, with the most famous example being radiotherapy. In the latter, high-energy electromagnetic fields destroy cells along linear paths and neutralize localized tumors. In contrast, infections spread very quickly over large regions or multiple tissues, creating a setting that is unsuitable for radiotherapy. The alternative, non-ionizing radiations tend to excite collective energy modes in the tissue, producing heating or vibrations instead of molecular, atomic or nuclear transitions, thus penetrating deeper in the tissue. In practice, this means the field is less invasive like ultrasound devices. However, low-energy radiations found no direct application to combat or disrupt ongoing viral infections. The development of effective and non-invasive methods for inactivating viruses is crucial for controlling the spread and future pandemics, as well as providing better patient care^8^.

Here, we demonstrated a direct application of low-intensity ultrasonic waves on the infectivity of the SARS-CoV-2 virus. The application was first predicted by Wierzbicki et al using numerical simulations and mechanical models, in which the authors proposed that ultrasonic waves could trigger resonances within the virus. In their first model, the authors found capsid and spike resonances between 100 and 500 MHz^3^. An improved model has been proposed to account specific regions in the spike protein^4^, specifically the *α*-helices and tropocollagen molecules, leading to resonances in the 1–20 MHz range, far closer to our own experimental findings. Having said that, we emphasize that our experiment alone cannot prove the model in Ref.^4^ correctly describes the underlying physical mechanisms resulting in the loss of viral infectivity.

Furthermore, our data suggests that the frequency band 5–10 MHz target a common viral structure across the variants considered here, whereas lower and higher frequencies are far less effective. Like other coronavirus, four structural proteins dictate the overall geometry of SARS-CoV-2, namely, N, S, E, M. The N-protein associates with the viral genome (around 30 kb linear single strand RNA) forming the viral core, being the most abundant protein^9^. E and M-proteins together with the glycoprotein spike S protein form the envelope. Mutations associated with the various strains affect either structural or non-structural proteins. These changes in turn can change physical properties of the virus, such as the Young modulus, a parameter that describes the rigidity of a medium and governs the medium ‘s wave velocity and acoustic impedance.

Our findings strongly suggest that ultrasonic acoustic waves have virucidal effects across several strains. SARS-CoV-2 particles in solution are 80–140 nm in diameter, 3–4 order of magnitude smaller than the typical wavelengths 0.1–5 mm used in medical devices. From a practical perspective, no meaningful interactions should be expected given the discrepancy in scales. However, the core-shell structure of the virus play a key role in the process. Recent studies in acoustic scattering show that internal resonances occur at the small particle limit, provided the scattering center is viscoelastic^10^. Under small forces, a viscoelastic material deforms and returns to its original form once the force ceases to be applied. Unlike purely elastic materials, a fraction of the energy dissipates due to internal resistances during the deformation. In that sense, SARS-CoV-2 fulfills the description of viscoelastic material as no plastic deformations were observed under the use of AFM probes. Indeed, the assemblies formed by viral envelope and nucleoproteins play the role of core-shell compounds and the scattering of acoustic waves could lead to acoustic resonances. This hypothesis is in agreement with our observations that the nucleoproteins are being expelled from inside out like a popcorn.

### Applications and safety

Our findings demonstrate standard ultrasound imaging devices can be used to neutralize viable SARS-CoV-2 particles in vitro. These equipment are properly regulated and deemed safe for human use, finding applications in imaging and diagnostics. Typical frequencies range from 1 to 40 MHz, with intensity *I*_0_ between 1 and 10 mW/cm^2^ at the transducer. Due to the attenuation law, the intensity of acoustic waves decay exponentially with penetration depth *z* according to the formula *I*(*z*) = *I*_0_e^−*μz*^. The characteristic length 1*/μ* depends on mechanical properties of the tissue and also on the propagation mode and frequency of the wave. Taken together, these parameters govern how far acoustic waves can travel throughout human tissues. As a rule of thumb, low frequencies tend to travel further, which opens opportunities for non-invasive therapies. In particular, it opens a venue to treat regions that are otherwise inaccessible such as the brain.

Moreover, recent studies indicate that no significant differences were observed in abnormal fetal ultrasound findings in pregnant persons who tested positive for SARS-CoV-2, indicating the safety of the equipment in humans^11^. The increase in temperature is associated with ultrasound exposure, and elevated temperature inhibits SARS-CoV-2 replication^12,13^. We did not observe any significant differences in the temperature of the culture medium during ultrasound exposure.

### Limitations

The primary focus of this research is the virucidal impact of ultrasonic waves. Given that the theoretical predictions for resonance frequencies fell within the 1–20 MHz range, we utilized standard ultrasound devices for imaging and diagnostics, as they align with the desired frequency range and are readily available. The in vitro experiments were conducted in a BSL-3 environment using three different strains. The analysis of confocal images and TCID50 assays were supplemented with AFM and SEM images. However, several limitations were identified in this approach.

Firstly, while ultrasound imaging devices are convenient to use and acquire, they lack precise controls for frequencies. In fact, the frequency distribution for each preset mode is designed to capture specific tissues, making them not easily adjustable. Furthermore, different modes may have overlapping ranges with different frequency distributions. These factors collectively impede our ability to accurately determine the resonance frequencies or quantify the changes in them due to mutations without external piezoelectric transducers.

Secondly, samples for AFM and SEM were prepared and dried overnight with cellular byproducts present in the stock medium. That is, no subsequent purification step was applied to the stock, control, and treated samples, as it was unknown at the time whether collisions in the ultracentrifuge would contribute to ultrasound resonances. However, the excess debris can deform the particles during the drying step, complicating the assessment of structural changes and the full characterization of the particle, even after washing steps. Coating the substrate with antibodies and performing AFM in solution could resolve this issue.

Finally, although the experiments were performed in absence of pre-installed bubbles, restricted to MI levels below the usual cavitation threshold MI ≈ 0.7, and Fig. 2A rules out cavitation as the main driver for the phenomenon, X-ray coupled with external piezoelectric sensors are needed to assess whether the formation of micro- or nano-bubbles is facilitated by the virus in solution (particulate) in vitro.

## IV. CONCLUSION

The global pandemic caused by SARS-CoV-2 has shown the urgency to develop new methods to combat viral infections, adding to the current known and well-tested methodologies such as vaccination and outbreak surveillance. Here, we investigate the potential virucidal effect of ultrasound medical devices as a non-invasive and broad-spectrum treatment against various strains of SARS-CoV-2. Our findings suggest that certain ultrasound frequencies can effectively nullify the virulence of SARS-CoV-2. More specifically, the 5–10 MHz band showed a lasting effect over the replication cycle of the virus across the strains considered here. This result suggests a shared resonance frequency – and thus conserved structure – that breaks the envelope apart. Typically, mismatching scales involving particles (80–140 nm) and waves (0.1–1.0 mm) produce no scattering events. However, theoretical predictions (see Discussion) suggest distinct mechanisms in which the wave transfer the energy to the particle, leading to resonances followed by envelope failure, in agreement with our experimental observations. Also, no significant temperature increase was observed during ultrasound exposure supporting of a mechanical energy transfer. Finally, due to the high-penetration power of low-frequency acoustic waves in animal tissue, our findings open up new options for the development of non-invasive antiviral treatments for spherical virus, particularly for regions that are of difficult access via traditional methods such.

## V. MATERIALS AND METHODS

### A. Virus stock production

The SARS-CoV-2 parental Wuhan, SARS CoV-2 gamma (P1), and SARS CoV-2 delta variants were used for in vitro experiments (accession code SRR22561695/PRJNA909758/SAMN32093250), under strict biosafety level 3 (BSL3) conditions at the Ribeirão Preto Medical school (Ribeirão Preto, Brazil). Briefly, viral inoculum (1:100 ratio) was added to the Vero E6 cells, and the culture was incubated (48 h, 37 °C, 5% CO2 humidified atmosphere) in DMEM without FBS but supplemented with antibiotic/antimycotic mix (Penicillin 10,000 U/mL; Streptomycin 10,000 *μ*g/mL; Sigma-Aldrich; cat. P4333) to optimize virus adsorption to the cells. After confirming the cytopathic effects of the viral replication over cell monolayer, cells were detached by scraping, harvested, and centrifuged (10000 ×g, 10 minutes, room temperature). The resulting supernatants were stored at -80 °C until use. SARS CoV-2 variants titration was assessed using standard limiting dilution to determine the 50% tissue culture infectious dose (TCID50).

### B. In vitro SARS-CoV-2 infection and US exposure

Vero E6 cells were infected with SARS-CoV-2 previous to the 30 minutes exposure to linear transducers with bandwidths 3–12, 5–10, or 6–18 MHz. The transducers were attached to ultrasound high-definition imaging devices, either MyLab 60 (Esaote) or Envisor HD (Philips). Cells were infected at a multiplicity of infection (MOI) of 1.0 with infectious clone SARS-CoV-2 or mock with infection media for 24 hours to evaluate the infection and replication process by immunofluorescence and confocal microscopy. The productive viral particle was assessed by TCID50 assay. The analysis was performed in technical triplicate. The culture medium temperature was measured using a thermal camera (FLIR One Pro, Flir).

### C. Immunostaining and confocal

For SARS-CoV-2 detection in vitro, Vero-E6 cells were plated in 24-well plates containing glass coverslips, fixed with PFA 4% at RT for 10 minutes, and blocked with 1% bovine serum albumin (BSA; Sigma-Aldrich; cat. A7906) and 22.52 mg/mL glycine (Sigma-Aldrich; cat. G8898) in PBST (Phosphate Buffer Saline + 0.1% Tween 20) at RT for 2 hours. The coverslips were stained with the following antibodies: rabbit anti-spike protein (Invitrogen; cat. 703959; 1:500) and mouse anti-dsRNA (J2; dsRNA, SCICONS English & Scientific Consulting Kft., clone J2-1909, cat.10010200; 1:1,000). After this, samples were washed in PBS and incubated with secondary antibodies: alpaca anti-mouse IgG AlexaFluor 488 (Jackson ImmunoReseacher; Cat. 615-545-214; 1:1,000) and alpaca anti-rabbit IgG AlexaFluor 594 (Jackson ImmunoReseacher; Cat. 611-585-215; 1:1,000). Slides were then mounted using Vectashield Antifade Mounting Medium with DAPI (Vector Laboratories; cat. H-1200-10). Images were acquired by Axio Observer combined with an LSM 780 confocal microscope (Carl Zeiss) at 630X magnification at the same setup of zoomed and laser rate. Images were acquired and analyzed using Fiji by Image J.

### D. Titration TCID50

To evaluate the effect of exposure to the US on SARS-CoV-2 infectivity, the virus stock was diluted 1:100 in each of the following: DMEM and/or US-exposed SARS-CoV-2. These two SARS-CoV-2 preparations were incubated for 1 min at room temperature, serially diluted 10-fold in DMEM, and then 100 *μ*L of each dilution was inoculated in quadruplicate monolayers to determine the virus titer by TCID50 in Vero CCL-81 cells in 96-well plates.

### E. Statistics

Statistical significance was determined by one-way ANOVA followed by Bonferroni ‘s post hoc test, *p* < 0.05 was considered statistically significant. Statistical analyses and graph plots were performed and built with GraphPad Prism 9.3.1 software.

### F. AFM and SEM images

For the AFM and SEM images, samples of the SARS-CoV-2 stock solution were exposed to UV light before frozen at -80 C, for use outside BSL-3. Poly-L-Lysine (Electron Microscopy Sciences; CAS 25988-63-0) was left to incubate for 5-10 minutes clean Si substrates (nFinitu; 5 mm x 5 mm diced silicon wafer). Inactivated samples were placed onto the coated substrate and fixed with 4% formaldehyde solution. After a washing cycle, the samples were dehydrated with a series of increasing concentration (40–100%) of ethanol. For SEM, the samples were sputtered with 10 nm thick graphite coat and observed in the Sigma Zeiss FE-SEM microscope (Carl Zeiss) at 2kV. For AFM, the samples were not coated and observed in an Icon-Dimension AFM (Bruker) without conductive coating, in tapping mode.

## ACKNOWLEDGMENTS

OMB would like to thank Daniela Gravina Stamato for the discussions on clinical ultrasonography and for the suggestions provided at the beginning of the research. Thanks are also extended to Tomasz Wierzbicki for the correspondence regarding the study of ultrasound resonance. The authors are grateful for the financial support provided by FAPESP: FPV 2020/07645-0, GN 2023/07241-5, FQC 2013/08216-2 and 2020/05601-6, EA 2019/26119-0, OMB 2018/22214-6 and 2021/08325-2. OMB also acknowledges the grant CNPq 307897/2018-4.

